# Predictive modeling of co-evolving growing populations

**DOI:** 10.1101/633750

**Authors:** Philipp M. Altrock, Meghan C. Ferrall-Fairbanks, Gregory J. Kimmel

**Affiliations:** Integrated Mathematical Oncology Department, H. Lee Moffitt Cancer Center and Research Institute, Tampa, FL 33612 USA

## Abstract

How can we best explore and fit density and frequency dependence in the evolutionary and ecological dynamics of growing tumors? Here, we present an introduction to recent developments in our lab, and give two examples of complex interaction-driven tumor growth combined with statistical outcomes of treatment.

## I. COMPETITIVE GROWTH DYNAMICS IN CANCER

Evolutionary game theory (EGT) allows to study the long-term success of co-evolving types that at least temporally coexist in a population, and whose interactions are quantified by a strategic interaction pattern, or game. The game is typically formulated as a table that ascribes payoffs to every pairwise interaction between types, e.g. cell types or strategies. Based on these interactions, EGT is able to mathematically predict the changes in the relative abundance of types, as payoffs are translated to proliferative advantage, or fitness. Of particular interest in this context is frequency-dependent selection, in which the fitness landscape changes as the population evolves^1^. This framework is useful to identify equilibria in which types may coexist and to make statements about the stability of these equilibria^2^, and to elucidate the rules of phenotype-driven changes and overall population diversity^3^. EGT traditionally describes populations with a predefined set of phenotypes, and the dynamics happen in populations of fixed size or of universal growth^4^. Only recently have a flexible number of phenotypes^5^ or a population that changes in size^6^ been considered in terms of their ability to generate and maintain coexistence of types under frequency dependent selection^7^. Recent developments show that EGT is an exciting field for mathematical modelling of cancer, yet proper integration of inhomogeneous growth models, i.e. density dependence, paired with frequency dependent selection is still lacking behind, but has wide applicative importance^8^. In this presentation we revise how individual-based processes that include frequency-dependence and density-dependence can be analyzed, and provide two examples based on recent data of cancer growth.

## II. INDIVIDUAL BASED MODELING

Let us assume that there are *m* cell types in the growing population, each growing with the net growth rate

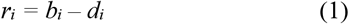

in mono-culture, where *b*_*i*_ and *d*_*i*_ are the intrinsic birth and death rates of each type, potentially themselves system dependent^8^. Next, we can assume that the payoff of type *i* interacting with type *j* is

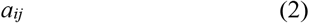

such that a deterministic growth model that emerges directly from an individual based (stochastic) formulation^9^ can be written as

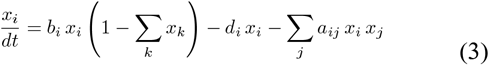

which incorporates the interactions (payoffs) as negatively affecting each cell type, resembling the Lotka-Volterra equation^6^. Here, we assume that the cellular densities are normalized with respect to the system’s carrying capacity, which leads to the logistic multiplier of the birth rates.

## III. FIBROBLAST-MACROPHAGE GAME IN LYMPHOMA

It is important to consider potential impacts of radiation therapy on lymphoma tumor microenvironment, and how those changes could help or hinder lymphoma cell growth. For example, irradiation can cause fibroblasts to be growth-arrested^10^ and induce a senescence-associated secretory phenotype, which may have tumor-suppressive function^11,12^. We modeled lymphoma cell growth with different combinations of wildtype and senescent fibroblasts, and in co-culture of activated or inactivated macrophages. The goal is to tease apart the effects of tumor-typical complex microenvironmental conditions on lymphoma cell growth. We used an ordinary differential equation model of the form of (3) to describe the birth, death, and interactions between the three cell types of fibroblasts (FIB), macrophages (MAC) and lymphoma (LYM). We iteratively estimated the rates from single, double, and triple culture experiments using cell counts over time. We tracked the changes in parameters due to the addition of each new cell type and extrapolated lymphoma growth with *in silico* simulations. In this system, the competitive interactions between the three types are summarized in the following pairwise-interaction payoff table:

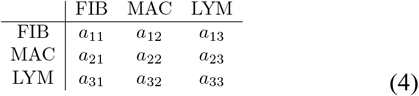

This table describes inter-species (*a*_*ij*_, *i*≠*j*) and intra-species (*a*_*ij*_, *i*=*j*) competition. In the deterministic limit of large population sizes, the rate equations for this triple culture system are described by Eq. (3). We found that senescence cell types, especially in combination with lymphoma cells and macrophages, predict an expansion of the tumor. Figure 1 shows the predict growth after 3 weeks as indicated by log-fold-change of the tumor cell population. The size of each point scales based on the log-fold-change; smaller points represent less growth and larger points represent more lymphoma growth. The absence of points in parameters space indicates a contracting lymphoma population. Simulations with both irradiated fibroblasts (IRR) and super-repressed/inactive macrophages (ASR) predicted the largest lymphoma population growth (pink points), even with 10-fold more fibroblast and macrophages initially present compared to lymphoma cells. Super-repressed macrophages with wildtype fibroblasts predicted some contraction of the lymphoma population, while irradiated fibroblasts in the presence of wildtype macrophages predicted unchecked lymphoma growth when low numbers of macrophages were present.

**Figure 1.**
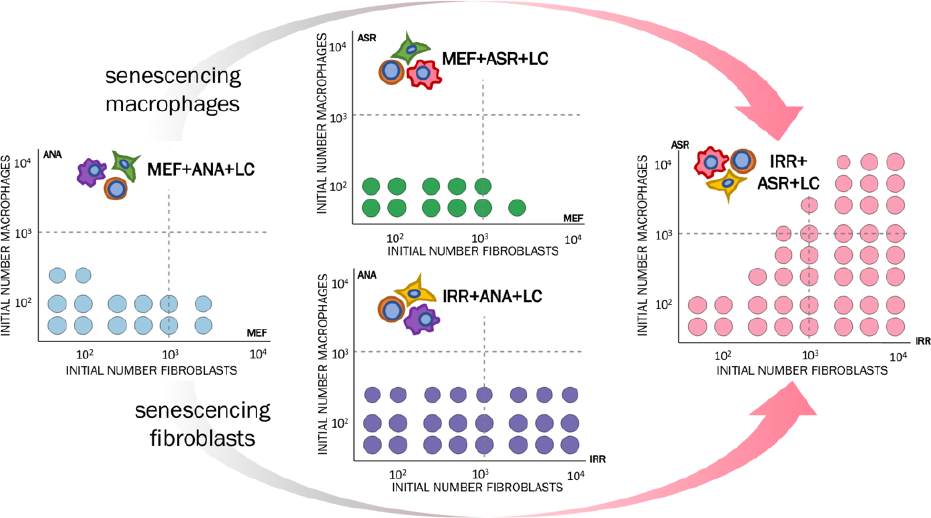
Forecasted lymphoma growth in different (micro-)environments. MEF=normal fibroblasts. IRR=Irradiated fibroblasts. ANA=normal macrophages. ASR=super-repressed macrophages. LC=lymphoma cells.

## IV. CAR T-WILDTYPE T-CELL INTERACTIONS

Modern immune therapeutic interventions, such as CAR T-cell therapy, rely on the expansion of genetically engineered T-cells that have been taken from the patient to be altered *ex vivo*. CAR T-cell therapy represents a huge medical breakthrough in cancer treatment^13^, but comes with a set of harsh side-effects. Little research has been made to model the dynamics of CAR T-cells mechanistically. The utility of a mechanistic model includes being able to determine efficacy of treatment on individual patients and optimal proportion of CAR T cell subtypes which maximize favorable treatment outcomes and minimize potential risks for side effects. We developed a mechanistic model, which describes the dynamics of CAR T cell dynamics *in vivo* (Figure 2), and identified three crucial states (equilibria of the co-evolutionary system) based on CAR T-cells interacting competitively with wildtype T cells; tumor eradication, stable tumor (coexistence), tumor growth irrespective of therapy. Possible equilibria (long-term stable states) of the (stochastic) dynamical system should correspond to the following clinical states observed in patients: No response, transient response, and long-term response (cure). Due to potentially low tumor burden, these dynamics are inevitably stochastic, their deterministoic (mean-field) analog should be of a form similar to the model in Equation (3), that is we assume that growth is logistically dampened, as well as impacted by competitive interactions.

**Figure 2.**
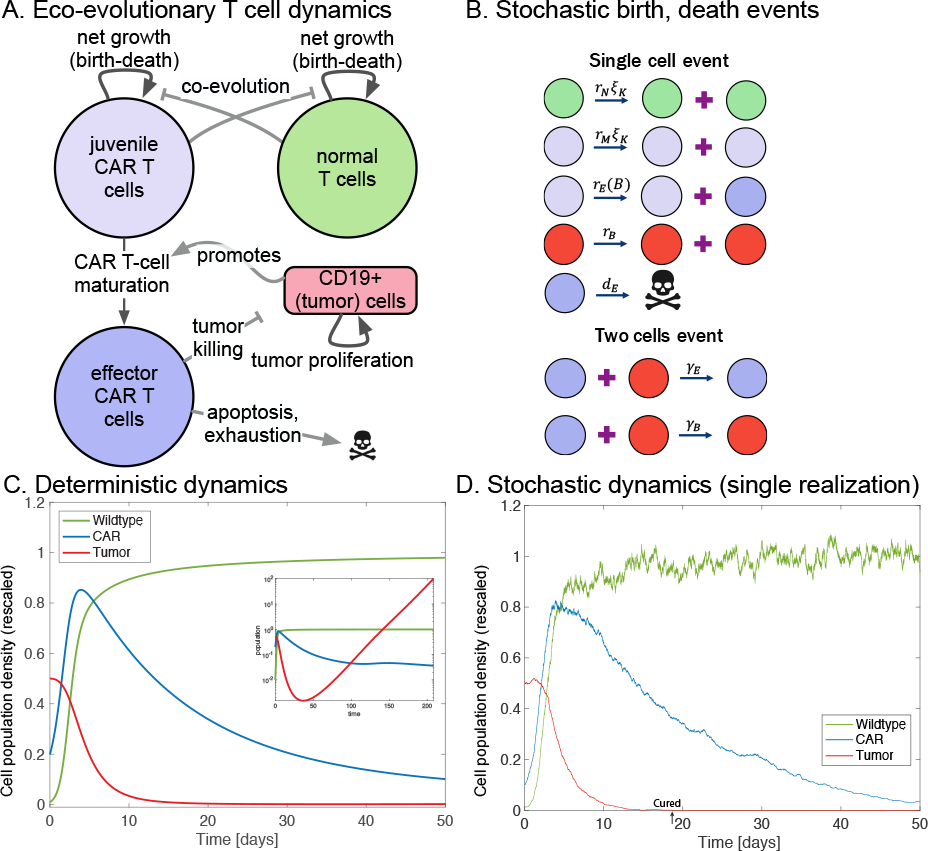
A: Model schematic (for an apppropriate non-linear ODE system). B: Stochastic model driven by birth, death and interaction events. C: Population dynamics of the CAR, wildtype and tumor towards a PD state. Initial treatment response is seen for 50 days followed by disease progression (inset). D: Stochastic population dynamics usig the same parameters.

## V. DATA FITTING

We assume that the system of interest has been measured over time, in multiple instances, such that longitudinal cell count data is available. Net growth, birth, and competition rates can be estimated from the experimental data in a step-wise fashion. First, the net growth rates can be estimated for each cell type assuming an exponential growth model from mono-culture experiments, as performed in previous work^14^. Then, individual birth and competition rates can be estimated from multi-culture systems, and L1 loss regularization-based machine learning can be used to determine interaction parameters payoffs, and potentially shrink the model structure. The number of parameters, is identified by varying the penalty coefficient and applying an Akaike information criterion (*AIC*) to determine the goodness of fit, using the sum of the residuals between the experimental data and the fitted estimates^15,16^. AIC estimates the quality of each model, given a collection of models for the data, which emerges for example by changing and simplifying the payoff table (4). Parameter estimation was performed in Julia using the DifferentialEquations, Optim, PenaltyFunctions, and Stats packages^17–19^.

